# Pupal cannibalism by worker honey bees contributes to the spread of Deformed wing virus

**DOI:** 10.1101/2020.11.25.396259

**Authors:** Francisco Posada-Florez, Zachary S. Lamas, David J. Hawthorne, Yanping Chen, Jay D. Evans, Eugene V. Ryabov

## Abstract

Transmission routes impact pathogen virulence and genetics, therefore comprehensive knowledge of these routes and their contribution to pathogen circulation is essential for understanding host-pathogen interactions and designing control strategies. Deformed wing virus (DWV), a principal viral pathogen of honey bees associated with increased honey bee mortality and colony losses, became highly virulent with the spread of its vector, the ectoparasitic mite *Varroa destructor*. Reproduction of *Varroa* mites occurs in capped brood cells and mite-infested pupae from these cells usually have high levels of DWV. The removal of mite-infested pupae by worker bees, *Varroa* Sensitive Hygiene (VSH), leads to cannibalization of pupae with high DWV loads, thereby offering an alternative route for virus transmission. We used genetically tagged DWV to investigate virus transmission to and between worker bees following pupal cannibalisation under experimental conditions. We demonstrated that cannibalization of DWV-infected pupae resulted in high levels of this virus in worker bees and that the acquired virus was then transmitted between bees via trophallaxis, allowing circulation of *Varroa*-vectored DWV variants without the mites. Despite the known benefits of hygienic behaviour, it is possible that higher levels of VSH activity may result in increased transmission of DWV via cannibalism and trophallaxis.

## Introduction

Pathogens, including viruses, exploit multiple transmission routes across different developmental stages which contribute to pathogen circulation and lead to diverse impacts on host physiology and life history^1^. Changes in modes of pathogen transmission could impose new evolutionary pressures on pathogens, in turn leading to pathogen phenotypic changes, including altered virulence^2^. Comprehensive models of transmission routes and their roles in pathogen circulation are essential for understanding pathogen evolutionary dynamics and development of control strategies.

Deformed wing virus (DWV)^3^, a principal viral pathogen of honey bees (*Apis mellifera*) associated with increased honey bee mortality and colony collapses^4–6^, has benefited from a novel transmission route in recent decades. Historically, DWV caused mainly covert infection characterized with low virus levels and transmission *via* food or individual bees^7^, but a dramatic increase of DWV virulence and infection levels was reported with the spread of ectoparasitic mite *Varroa destructor*. The mite feeds on the hemolymph and fat body tissues of pupae and adult bees and is an effective vector for viruses, including DWV^8–10^. *Varroa*-mediated transmission of DWV by direct injection into the insect hemolymph, allowing the efficient movement of viruses from infected bees to others, favored more virulent DWV strains. Genetic changes in DWV which occurred as a result of *Varroa* vectoring included reduction of genetic diversity and selection of particular strains^11–15^. Reproduction of *Varroa* mites occurs exclusively in capped honey bee brood cells, with the mite and the mite-infested pupae showing high levels of DWV^7,9^. Bees can suppress *Varroa* mite reproduction by selecting and uncapping *Varroa*-infested brood cells and removing infested pupae by co-called *Varroa* Sensitive Hygiene (VSH)^16–18^. It has been suggested that VSH could be accompanied by cannibalization of mite-exposed pupae^19^.

Cannibalism, consuming of conspecific individuals, occurs in many animals^20,21^. It is common in the eusocial Hymenoptera, ants^22,23^, wasps^24^, bees^25–27^ and termites^28,29^ throughout the growth and development of the social organization and may occur for a variety of reasons including the nutrient shortages, disease and pest outbreaks, environmental stressors, and colony disturbance^30^.

In honey bees, cannibalism is an essential part of social organization and colony-level hygiene is exercised through ecological, physiological, genetic and sanitary stressors^20,30–32^. Any developmental stages and castes can be cannibalized including developing queens. Honey bees show natural cannibalism behavior when workers police to control worker-laid eggs^33^ and remove diploid drone larvae^34^. Cannibalization of eggs or younger larvae can be stimulated by environmental conditions, unbalanced nutrition such as scarcity of pollen^25,27,30,35^, and when honey bees perform hygienic behaviors^17,18^.

Among the main risks associated with cannibalism is the increased spread of pathogens, in particular in the case of group cannibalism, *i.e.* when the prey is shared across a social group^36^. In invertebrates, ingestion of infected conspecific tissues is recognized as a route of virus transmission in insects and shrimp^37,38^. At the same time, reduction of the numbers of infected individuals by cannibalism might limit the spread of disease^21^. Although it was suggested that worker bees could be infected with DWV as a result of cannibalization of virus-infected bees^39^, this has not been experimentally investigated. One reason complicating the study of the impact of cannibalism on DWV circulation is the difficulty in distinguishing between DWV infection initiated by cannibalization and by other routes. To solve this problem we used genetically-tagged DWV carrying unique genetic markers, the green fluorescent protein (GFP) gene and an introduced unique restriction site^40^, allowing us to trace transmission of the virus. We also carried out pupal cannibalism experiments in controlled laboratory conditions rather than hives, thereby minimizing virus transmission from other sources. This study provides the first direct experimental evidence that cannibalization of pupae with high levels of DWV leads to infection in worker honey bees, and that DWV could then be shared extensively among worker bees by trophallaxis. Our results suggest that cannibalization of pupae infected with DWV by *Varroa* mites, removed as a result of VSH activity, could provide an efficient additional route for transmission of DWV, impacting virus circulation and virulence.

## Results

### High levels of DWV in partially cannibalized honey bee pupae removed by hygienic activity

Partially cannibalized pupae (n = 15) showing different degrees of damage, ranging from partially to completely removed heads, were collected from hygienically open cells of four colonies showing cannibalism by worker honey bees (Fig. 1a). In two of these colonies, some pupae (n=7) were sourced from hygienically opened brood cells containing *Varroa* mites at the time of collection. Notably, *Varroa* mites were found more often in hygienically opened brood cells containing partially cannibalized pupae than in capped brood cells (Colony #10: for partially cannibalized pupae 6 mite-infested and 1 mite-free, for capped cells 4 mite-infested and 84 mite-free, P = 0.0000016 Chi-square test for contingency table analysis; Colony #11: for partially cannibalized pupae 1 mite-infested and 2 mite-free, for capped cells 0 mite infested and 85 mite-free, P = 0.03409 Chi-square test for contingency table analysis), suggesting that these pupal cells were opened as a result of *Varroa* sensitive hygienic (VSH) behaviour. We also collected control pupae (n = 9) from capped *Varroa*-free cells from areas of the brood frames where the partially cannibalized pupae were sourced. The pupae of both the control and damaged groups were at white to pink-eye developmental stages. Quantification of DWV RNA by RT-qPCR (Fig. 1b, Supplementary Table 1) showed that the levels of DWV in the partially cannibalized pupae (range: 5.05 to 10.50 log_10_ GE/pupa; 7.39 ± 1.589 log_10_ genome equivalents (GE)/pupa, mean ± SD) were significantly higher than in the capped *Varroa*-free pupae (5.39 to 6.86 log_10_ GE/pupa; 6.05 ± 0.473 log_10_ GE/pupa, mean ± SD), P = 0.022, df = 23, ANOVA (Fig. 1b). There was no significant difference between these groups in the levels of honey bee actin mRNA (P = 0.560, df = 23, ANOVA) (Supplementary Table 1) confirming that no tissue degradation, potentially affecting RNA quality or actin expression, took place in the damaged pupae.

**Figure 1.**
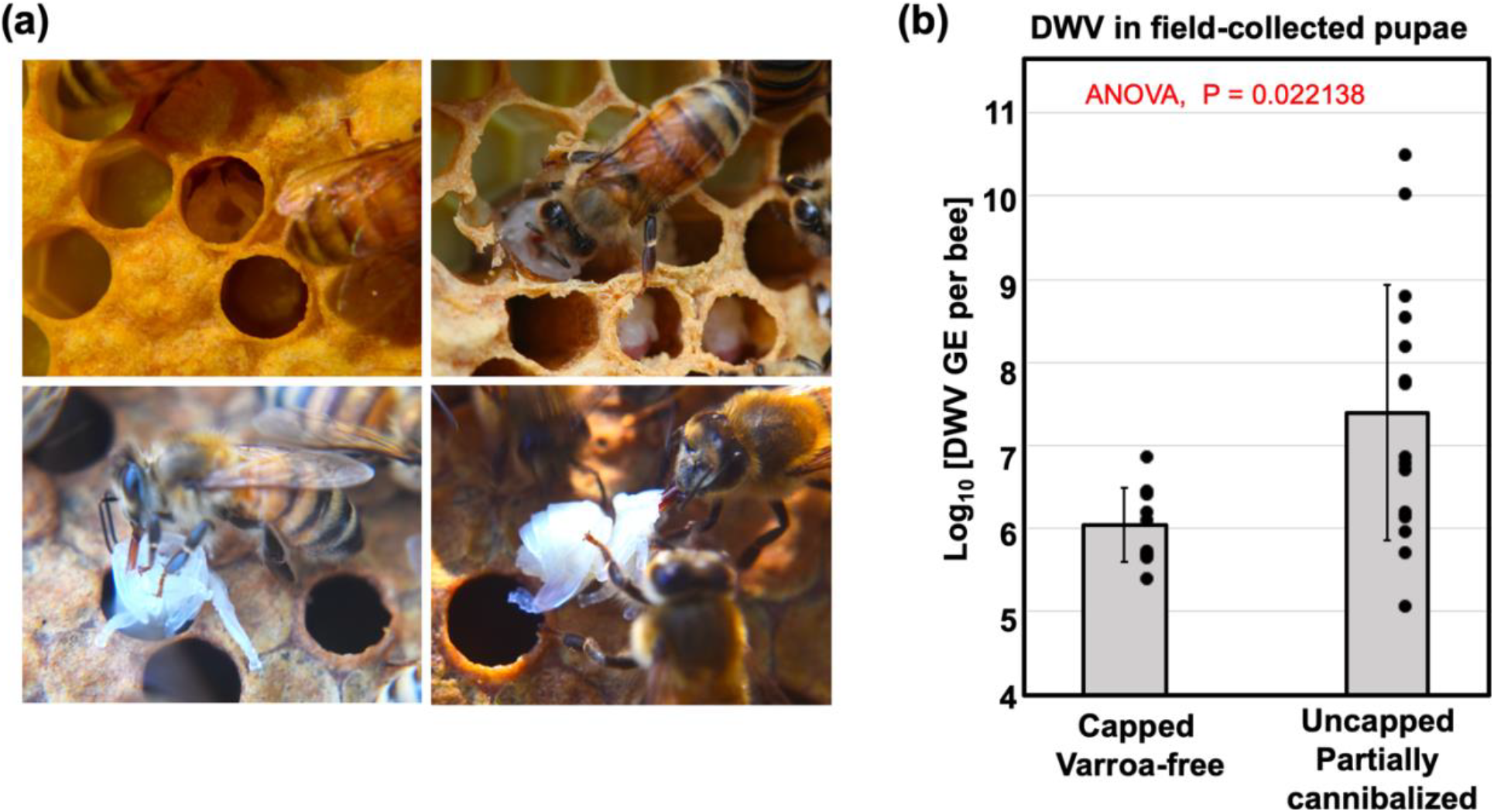
**(a)** Honey bee pupal cannibalization by worker bees. **(b)** Average DWV RNA loads in field-collected capped and in partially cannibalized uncapped honeybee pupae, with error bars indicating standard deviation. DWV copy number in individual pupae are indicated by black dots.

### Acquisition of DWV by worker bees as a result of cannibalism of pupae infected by *Varroa*

Experiment A tested if cannibalization of the pupae infected with DWV by *Varroa* mites could result in development of the virus infection in worker bees. It included injecting honey bee pupae with a filtered tissue extract containing DWV-GFP particles40, or by a phosphate buffer saline (PBS) control. After 48 hours (hr), when GFP fluorescence had developed in the DWV-GFP-injected pupae indicating virus infection (Fig. 2a), *Varroa* mites were placed on the pupae and reared for 72 hr to acquire the virus41. Then, these mites were transferred to new white-eye pupae (Fig. 2b, Pupa 2) which were reared for 5 days to allow transmission of the virus from *Varroa* and development of infections in the recipient pupae. Pupae from both PBS and DWV-GFP groups were cut sagittally, one half was used to extract RNA for molecular analysis of the DWV and GFP loads (Fig. 2c), and another half was offered to a group of 20 worker bees 4 days post-emergence (Fig. 2b). A control group of 20 worker bees received no pupae for cannibalization. Nearly complete cannibalization of “PBS” and “DWV-GFP” pupal tissues was observed after 12 hr incubation. The worker bees were maintained for an additional 10 days before sampling for molecular analysis of virus loads (Fig. 2b).

**Figure 2.**
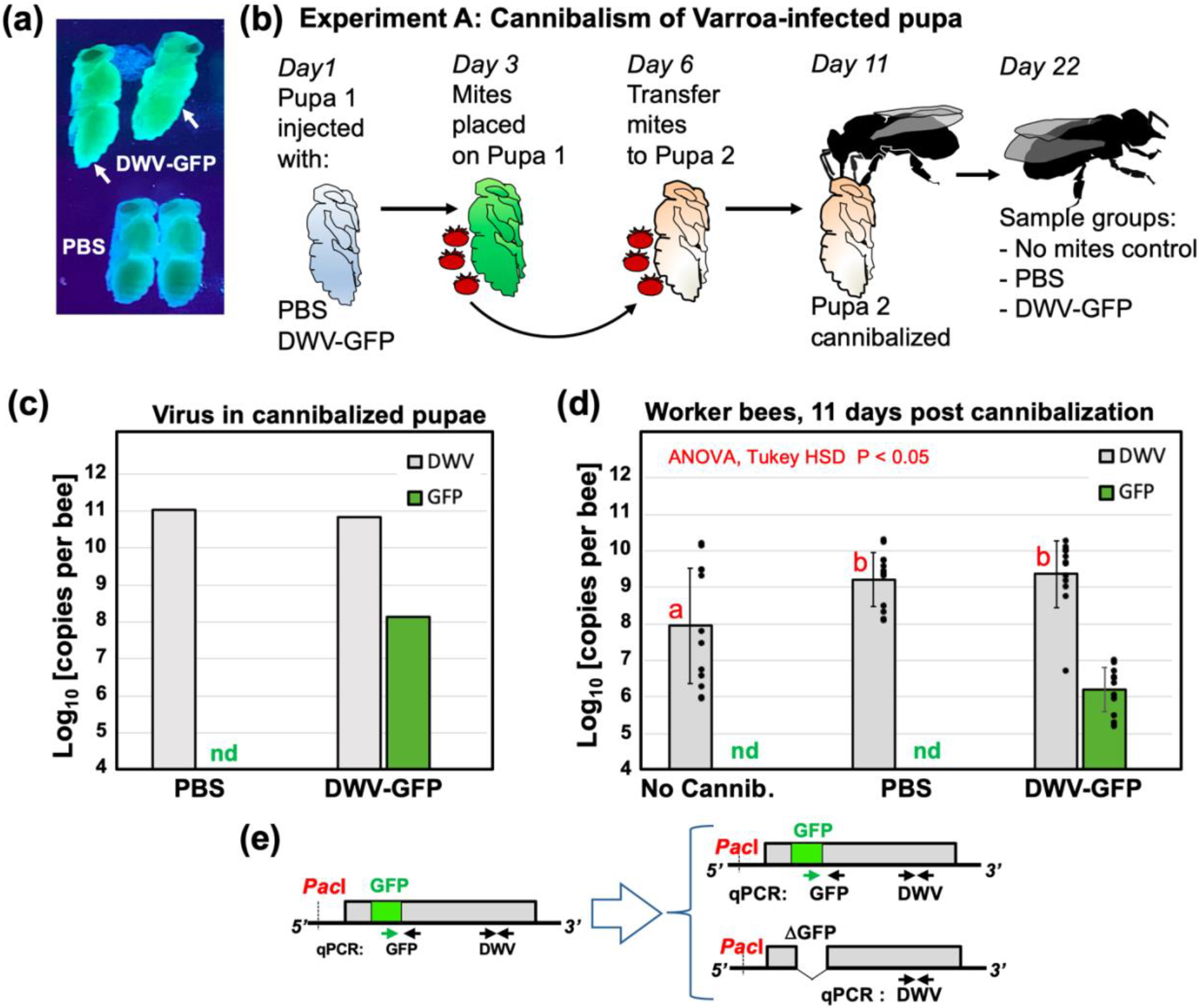
DWV infection in worker bees following cannibalization of pupae infected by *Varroa* mites (Experiment A). **(a)** Honey bee pupae, control and DWV-GFP-infected (pointed with arrows) which were used to rear Varroa mites (Pupae 1), illuminated with 395 nm UV light; **(b)** Schematic representation of the experiment. **(c)** DWV and GFP RNA loads in *Varroa* exposed Pupae 2 offered for cannibalization. **(d)** Average DWV and GFP RNA loads in individual worker bees, error bars indicate standard deviation. For DWV, red letters above bars indicate significantly and non-significantly different groups (ANOVA). DWV and GFP copy number in individual pupae are indicated by black dots, nd – not detectable levels. **(e)** Schematic representation of the DWV-GFP RNA genome and genetic changes following deletion of the GFP-coding sequence, positions of qPCR primers used for quantification of DWV and GFP RNA and genetic changes in DWV-GFP following deletion of the GFP-coding sequence are indicated.

RT-qPCR analysis of pupae which were exposed to *Varroa* mites for 5 days revealed that both “PBS” and “DWV-GFP” pupae had high levels of DWV RNA (Fig. 2c) (11.025 log_10_ GE/pupa, and 10.829 log_10_ GE/pupa, correspondingly). GFP RNA was detected only in pupae (8.1332 log_10_ GE/pupa) which were exposed to the mites which acquired DWV-GFP (Fig. 2d). The observed 496-fold (2.695 log_10_) excess of DWV over GFP in Pupa #2, which received DWV-GFP by mite transmission, could be a result of both accumulation of clone-derived DWV genomes with deletion of the GFP-coding sequence^40^ and transmission of wild-type DWV by *Varroa* mites. Indeed, high DWV levels in Pupa 2 of “PBS treatment” (Fig. 2c) suggested transmission of wild-type DWV by the *Varroa* mites used in this experiment.

Analysis of virus levels in worker bees 11 days post cannibalization (dpc) showed that “PBS” and “DWV-GFP” groups had similar levels of DWV (9.210 ± 0.7379 log_10_ GE/bee, and 9.3.54 ± 0.9149 log_10_ GE/bee, mean ± SD, correspondingly). DWV load in worker bees of the control group, which did not cannibalised pupa, “No Cannib.”, 7.940 ± 1.580 log_10_ GE/bee, mean ± SD, were significantly lower than in the groups which consumed DWV infected pupal tissue (P < 0.05, df = 35, ANOVA) (Fig. 2d, Supplementary Table 1). GFP RNA was present in the “DWV-GFP” worker bees at the levels of 6.196 ± 0.6051 log_10_ GE/bee, mean ± SD, which was lower than in the cannibalised “DWV-GFP” Pupa 2. The observation of an average 2502-fold (CI95 1030 to 6081-fold) excess of DWV over GFP in worker bees in the “DWV-GFP” treatment group could be explained by further loss of the GFP-coding sequence from the clone-derived DWV-GFP (Fig. 2e) and also by the presence of wild-type DWV.

### Trophallactic transmission of the virus acquired by pupal cannibalism

Experient A (Fig. 2) demonstrated that pupae infected with DWV by *Varroa* mites could be act as a source of infection when cannibalized by worker bees. The levels of DWV in these pupae, 10^10^ - 10^11^ GE (Fig. 2c), were similar to those observed in the pupae infected with DWV-GFP by injection^40^. Therefore, such injection-infected pupae could be used as an adequate replacement for *Varroa-*infected pupae in cannibalism experiments.

The impact of pathogens acquired by cannibalization depends on the number of individuals involved in cannibalism, either directly or through sharing the infected tissues^36^. Worker honey bees often exchange food by trophallaxis, which could allow the virus from the cannibalized tissues spread to a large number of workers. To test if such transmission takes place we devised an Experiment B (Fig. 3a) to investigate transmission of the infection between groups of worker bees separated by a wire mesh, allowing trophallactic contacts but not bee movement (Fig. 3b). A white-eyed pupa injected with DWV-GFP inoculum (7 log_10_ GE), which showed GFP fluorescence consistent with 10^10^ to 10^11^ GE of the virus 48 hr after injection (hpi), was divided into 5 equal parts, which were offered to 5 groups of 25 worker bees in the donor (cannibal) chambers of the cages. Controls, 5 groups of 25 worker bees, did not receive pupal tissue. Both control and experimental worker bees were 4 days old and were sourced from colony #2 with 0.5% *Varroa* mite infestation rate. Complete or nearly complete cannibalization of the offered pupal tissues was observed in each of 5 experimental cages. Five days later, newly emerged worker bees were placed into the “Recipient” chambers of all 10 cages (Fig. 3b) and were maintained for additional 10 days before sampling. The donor and recipient cages contained sugar syrup feeders, but to promote trophallaxis from the donor cage workers, the feeders were removed from the recipient cages for 8 hours during the first three days after placement the recipient group.

**Figure 3.**
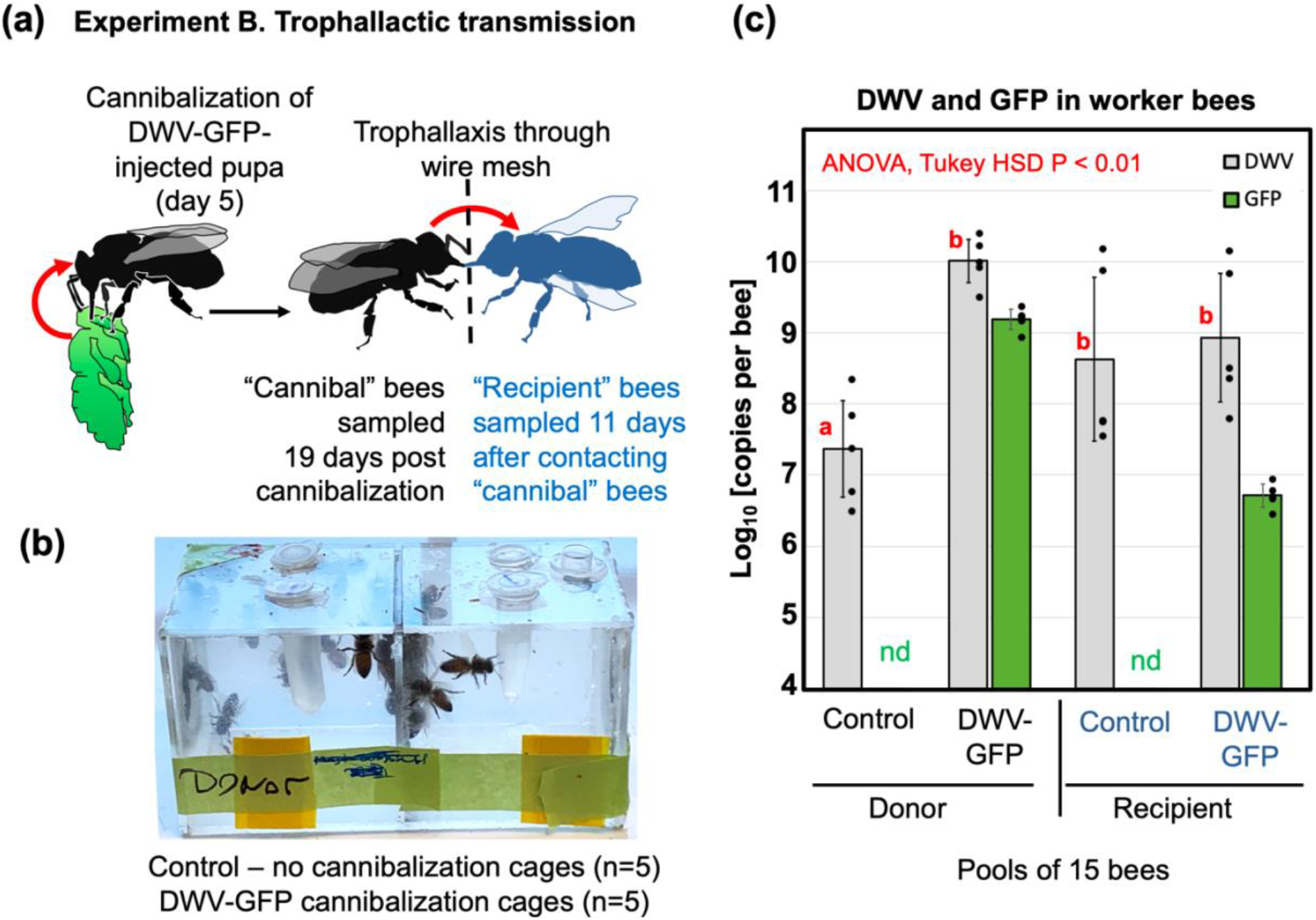
Trophallactic transmission of cannibalized DWV (Experiment B)**. (a)** Schematic representation of the experiment. **(b)** Design of the trophallactic cages. **(c)** Average per insect levels of DWV and GFP RNA in the pools of worker bees, error bars indicate standard deviation. For DWV, red letters above bars indicate significantly and non-significantly different groups (ANOVA). Average DWV and GFP copy number in the cage pools are indicated by black dots, nd – not detectable levels.

Average per-bee loads of DWV and GFP RNA were quantified by RT-qPCR in pools of 15 worker bees which were sampled from each donor chamber 19 days post cannibalism, and from each recipient chamber 11 days after contacting donor bees (Fig. 3c). Overall DWV levels, which included wild-type DWV, DWV-GFP and GFP deletion variants of this virus, were not significantly different in both recipient groups (“Control” and “DWV-GFP”) and in the donor “DWV-GFP” group (8.630 ± 1.1514 log_10_ GE/bee, 8.933 ± 0.904017169 log_10_ GE/bee, and 10.010 ± 0.3034 log_10_ GE/bee, mean ± SD, respectively) while DWV levels were significantly lower (p < 0.01, ANOVA) in the donor “control” group, 7.369 ± 0.6789 log_10_ GE/bee (Fig. 3c, Supplementary Table 1). The presence of GFP in the bees of both “Donor - DWV-GFP” and “Recipient - DWV-GFP” groups (9.190 ± 0.1416 log_10_ GE/bee and 6.712 ± 0.1635 log_10_ GE/bee, mean ± SD, respectively) but not in the control bees confirmed development of DWV-GFP infection following cannibalization of infected pupal tissues and further transmission of the tagged virus via trophallaxis (Fig. 3c, Supplementary Table 1). High levels of DWV in both control recipient groups was likely a result of a wild-type DWV infection, which was present in the recipient bees. Such contamination with wild type virus was not surprising because DWV is widespread in Maryland colonies^14^ and it is known that newly emerged worker honey bees may develop DWV infection even without receiving additional virus inoculum ^42^. Although the levels of GFP were approximately 7 times lower than those of DWV in the donor “DWV-GFP” groups (Fig. 3c), the *Pac*I restriction analysis of the RT-PCR fragment for the 5’ terminal region showed that all DWV present in these bees derived from DWV-GFP (Fig. 4a, lanes “Ex-B-Donor”). In the recipient “DWV-GFP” group, DWV levels were 167-fold higher than those of GFP, indicating that no more than 0.6% of population contained intact DWV-GFP (Fig. 3c). At the same time, *Pac*I digestion test which targeted the clone-derived DWV (Fig. 2e) showed that 42 % of the virus in the recipient “DWV-GFP” group derived from DWV-GFP (Fig. 4a, lanes “Exp-B-Recipient”).

**Figure 4.**
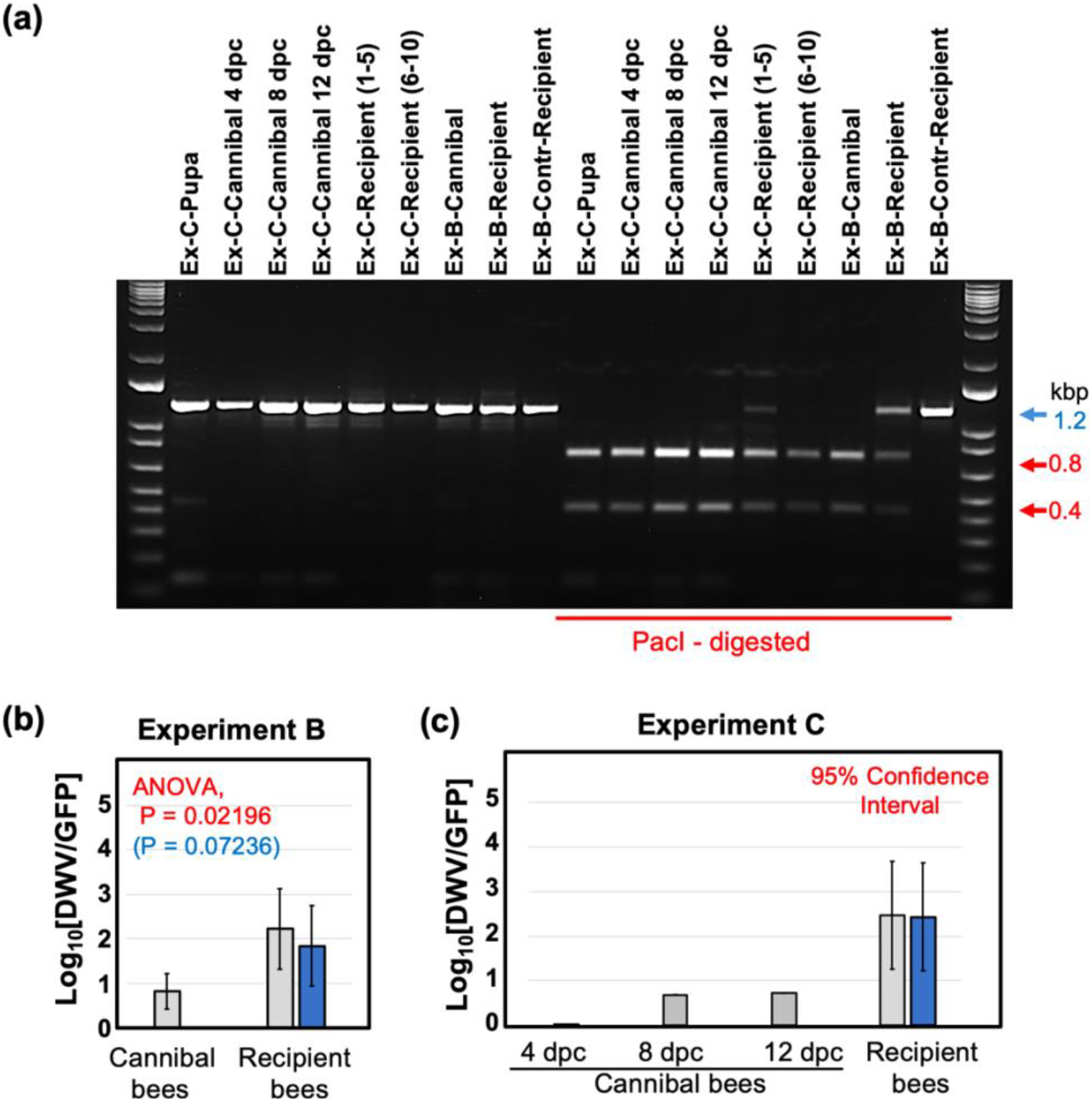
Dynamics of DWV-GFP in worker bees following cannibalism and trophallactic transmission. **(a)** Identification of DWV-GFP-derived viral progeny in the treatment group pools of Experiments B and C. The analysis included amplification of a 1237 nt RT-PCR fragments corresponding to the 5’-terminal region of DWV genome, digestion with *PacI*, and separation of the digestion reaction products by agarose gel electrophoresis. The untreated 1237 nt fragments (left) and *Pac*I-digested (right). The digestion fragments (left) derived from DWV-GFP, expected fragment sizes, undigested (blue arrow) and digested (red arrows), are shown on the right. Treatment groups are shown above, prefixes “Ex-B-” and “Ex-C-“ indicate samples of the Experiment B and C treatment groups, respectively. Two pools of 5 Recipient cages were analyzed for Experiment C. **(b, c)** Accumulation of GFP deletion variants derived from the DWV-GFP genome in the recipient bees which received the virus by trophallaxis from “Cannibal” bees. Columns indicate ratios between DWV RNA load and GFP RNA load in a sample, grey columns - for overall DWV levels, blue columns - for DWV originated from DWV-GFP (when wild-type DWV without *Pac*I site was present). Error bars indicate **(b)** standard deviation or **(c)**95 % Confidence Interval, for **(b)** ANOVA P-values for uncorrected DWV load (red) and corrected DWV load (blue) are shown.

DWV to GFP ratios, an indicator of the GFP loss from DWV-GFP, were increased in the recipient group compared to the donor group (Fig. 3c, 4b). This change was estimated as 25-fold for overall DWV, or 10-fold when only DWV-GFP-derived virus containing *Pac*I was considered and was statistically significant (P = 0.02196 for overall DWV levels, P = 0.07236 for the DWV-GFP-derived alone).

### Dynamics of DWV infection in worker bees following pupal cannibalism

Experiment C further investigated replication of DWV in worker bees at the individual insect level following consumption of DWV-infected pupae and further transmission of infection between worker bees when full contact was possible, similar to natural interactions between worker bees in the hive (Fig. 5a). This experiment involved three groups of 200 four-day old worker bees. The treatments included: no cannibalization control (Groups C), cannibalization of a *Varroa*-free pink eye pupae from the colony #2 injected with PBS with low levels of wild-type DWV, 95% Confidence Interval (CI) 5.353 - 5.753 log_10_ GE/pupa (Group T1), and cannibalization of the DWV-GFP injected pupa with high levels of the virus (95% CI: 10.702 – 11.027 log_10_ GE/pupa; Group T2). DWV and levels of GFP were determined by RT-qPCR in individual bees 4, 8 and 12 days post cannibalization, 10 insects were collected from each group for every sampling event. To investigate trophallactic transmission, 10 groups of 10 bees from each treatment group were collected at 4 dpc, marked and placed to the cages containing 90 newly emerged bees and reared together for additional 8 days. Then, the groups of 50 unmarked bees were collected and pooled for each of 30 cages and the levels of DWV and GFP were quantified. Molecular analysis showed that cannibalization of a *Varroa*-free honey bee pupa by the T1 bees did not result in development of a high-level virus infection in worker bees, and remained the same as in the control group which did not cannibalize (C) and as in the bees at the start of the experiment (T0), Fig. 3b. No GFP RNA was detected in the bees of T0, C and T1 of the “Cannibal” groups at any timepoint (Fig. 5b) and in C and T1 “Recipient” pools (Fig. 5c) (Fig. 5b). Cannibalisation of DWV-GFP infected pupae resulted in the development of DWV-GFP infection in worker bees (Fig. 5b, groups T2). At 4 dpc, the levels of DWV in T3 group were significantly higher than in the control group C and in the T1 groups which cannibalised non-injected pupa (Fig. 5b), ranging from 6.368 to 8.165 log_10_ GE/bee (7.627 ± 0.4614 log_10_ GE/bee, mean ± SD). Similarly high levels of GFP RNA were observed in 9 out of 10 bees, reaching 8.153 log_10_ GE/bee (7.368 ± 1.1453 log_10_ GE/bee, mean ± SD), with a single bee in this group having undetectable levels of GFP and DWV loads similar to those in bees of groups C and T1, 6.368 log_10_ GE/bee. Such nearly uniform distribution of DWV-GFP among 200 bees in the T2 group suggests that a high proportion of bees was involved in cannibalism and/or sharing of the virus-infected pupal tissue by trophallaxis. DWV-GFP infection continued to develop in T2, bees, exceeding 10^9^ copies per worker at 8 dpc (4 out of 10 sampled bees, highest level 10.620 log_10_ GE/bee, 8.798 ± 1.1080 log_10_ GE/bee, mean ± SD), and maintaining these levels at 12 dpc (with 2 out of 10 sampled bees, highest level 10.632 log_10_ GE/bee, (8.360 ± 1.0770 log_10_ GE/bee, mean ± SD) (Fig. 5c). The GFP RNA loads in “Cannibal T2” groups at 8 and 12 dpc (8.509 ± 0.735493769 log_10_ GE/bee and 8.137 ± 0.7905 log_10_ GE/bee mean ± SD) in these bees were slightly lower than those of DWV RNA (Fig. 5b). However, it was demonstrated by the complete digestion of cDNA fragments corresponding to the 5’ regions of DWV RNA with *Pac*I demonstrated that DWV, which did not carry the GFP insert, derived from DWV-GFP (Fig. 4a, lines “Ex-C-Cannibal-4, −8, −12 dpc”).

**Figure 5.**
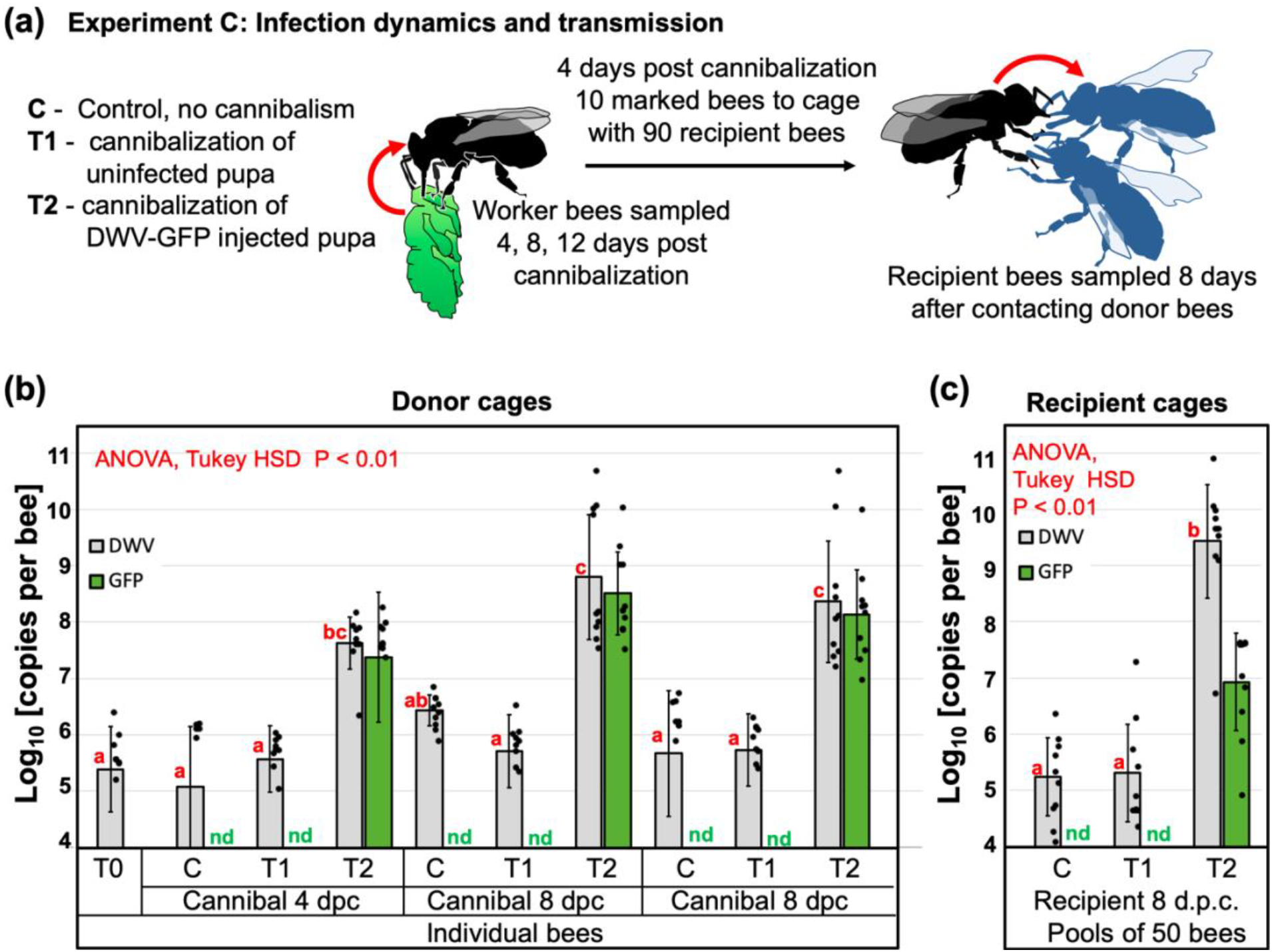
DWV dynamics in worker bees following cannibalization (Experiment C). **(a)** Schematic representation of the experiment. **(b)** Average DWV and GFP RNA loads in individual worker bees of the “cannibal” group, error bars indicate standard deviation. For DWV, red letters above bars indicate significantly and non-significantly different groups (ANOVA). **(c)** Average DWV and GFP RNA loads in the pools of worker bees of the recipient groups, error bars indicate standard deviation. For DWV, red letters above bars indicate significantly and non-significantly different groups (ANOVA). DWV and GFP copy number in individual pupae are indicated by black dots, nd – not detectable levels.

Experient C also tested the ability of bees which acquired DWV-GFP by cannibalization to transmit the virus to naïve worker bees when they are reared together. This was done by collecting 10 groups of 10 marked bees from each of three Donor cages at the 4 days after cannibalisation. The worker bees were marked and then transferred to Recipient cages containing 90 naïve newly emerged worker bees after which recipient bees were reared for 8 days (Fig. 5a). Molecular analysis of DWV and GFP RNA loads was carried out for the pool of 50 recipient unmarked bees for each of 30 Recipient cages (10 for each of 3 groups). The highest levels of DWV (9.427 ± 1.009 log_10_ GE/bee, mean ± SD, range 6.753 to 10.842 log_10_ GE/bee) were observed in the cages of the T2 group which received bees which cannibalized DWV-GFP infected pupa. Levels of DWV in T2 Recipient cages were significantly higher (P < 0.01, ANOVA, Tukey ‘s HSD) than those of the Control C group which did not cannibalise pupae (average 5.2414 ± 0.6944 log_10_ GE/bee, mean ± SD, range 4.1511 to 6.3929 log_10_ GE/bee) or T1 group, which cannibalised *Varroa*-free pupa (average 5.3078 ± 0.8668 log_10_ GE/bee, mean ± SD, range 4.7129 to 7. 2896 log10 GE/bee) (Fig. 5c). GFP targets were detected exclusively in T2 group cages, 6.930752384 ± 0.865399473 log10 GE/bee, mean ± SD, range 4.976 to 7.641 log10 GE/bee (Fig. 5c), indicating transmission of DWV-GFP acquired by cannibalism. Analysis of the RT-PCR fragment corresponding the 5’ region of DWV populations from group T2 showed that 93 % of DWV contained the *Pac*I restriction site unique for the cDNA clone-derived virus, indicating that a majority of the virus population originated from DWV-GFP. At the same time, RT-qPCR showed that DWV to GFP ratios in Recipient T2 groups were approximately 290:1 (2.462 log_10_), (or 268:1 (2.428 log_10_) if only clone-derived virus was considered. The DWV to GFP ratios in the T2 Cannibal bees were well below the 95% confidence limit for DWV to GFP ratios for T2 Recipient cages (Fig. 4c). In the Cannibal T2 group, nearly equimolar levels of DWV and GFP were observed at 4 dpc. As infection developed, accumulation of the viral variants with the deletion of GFP-coding sequence resulted in increase of the DWV to GFP ratios at 8 dpc and 12 dpc to 4.5 and 5.2, respectively (to 0.688 log_10_ and 0.720 log_10_, respectively) (Fig. 4c).

## Discussion

High-throughput sequencing has allowed the comprehensive characterization of invertebrate viromes, allowing the discovery of many novel viruses43, but understanding virus biology, including transmission routes, is lagging behind. This study investigated impacts of cannibalization of pupae by adult worker bees on circulation of DWV, the principal viral pathogen of honey bees3. Pupae cannibalized in *Varroa*-infested colonies were likely to be uncapped as a result of *Varroa* sensitive hygienic activity (Fig. 1), and some of these partially cannibalized pupae were shown to contain high levels of DWV consistent with overt DWV infections (Fig. 1b).

Testing the spread of DWV in colonies has proved to be difficult because it was not previously possible to distinguish between virus acquired by worker bees *via* cannibalism or *via* other routes, considering the nearly ubiquitous spread of DWV. Therefore, the role of cannibalism in maintenance of DWV infection has remained speculative so far. To test this hypothesis, we investigated cannibalism and transmission in controlled experimental conditions, using genetically tagged DWV isolate that allowed us to trace infections. This tagged virus containing a GFP insert^40^ was based on the cDNA clone of a virulent DWV isolate originated from *Varroa*-infested pupae sourced from a dying colony^15^, therefore this variant is suitable to study transmission of DWV acquired as a result of hygienic removal and cannibalization of *Varroa*-infested pupae.

We demonstrated that cannibalization of honey bee pupae infected with DWV either by *Varroa* mites (Fig. 2) or artificially infected with this virus by injection (Figs. 3 and 5), which contained high levels of the virus (95% CI: 10.702 – 11.027 log_10_ GE per pupa), resulted in infection levels typical for overt DWV infection, above 9 log_10_ GE, reaching 10.842 log_10_ GE, in worker bees at 8 dpc (Figs. 3 and 5). The levels of DWV-GFP in the pupae, cannibalization of which resulted in development of infections in workers, (Fig. 2c) were similar to those in some partially cannibalized pupae which were uncapped in hives as a result of VSH activity (Fig. 1) indicating that infection of workers as a result of pupal consumption could take place under natural hive conditions. Therefore virus-infected cannibalized pupae could act as “superspreaders” infecting large number of worker bees. For example, Experiment C showed that after cannibalization of a single pupa in a cage with 200 worker bees, 24 to 148 bees (CI95 12.2 – 73.8%) had developed overt DWV levels (Fig. 5b). At the same time, Experiment C showed that cannibalization of pupae with low levels of DWV, typical for covert infections, with 95% CI 5.353 – 5.753 log_10_ GE/pupa, did not result in development of overt level infection in worker bees (Fig. 5c, Group T1). It is possible that honey bees are adapted to suppress development of infection when tissues with low DWV are acquired orally^14^, but are unable to resist infection when high doses are ingested via cannibalized pupal tissue. Pupae with high DWV levels (above 9 log10 GE) are associated with *Varroa* mite infestation. Considering that *Varroa* became a parasite of *A. mellifera* very recently, it is possible that *A. mellifera* did not evolve antiviral defenses that might allow them to withstand high viral doses orally.

Considering that nurse worker honey bees are actively exchanging consumed food from mouth to mouth by trophallactic interactions^44^, such transmission of DWV acquired by cannibalism was tested in Experiments B and C (Figures 3, 5). It was demonstrated that the virus was readily transmitted by trophallaxis from cannibalistic bees to naïve recipient worker bees, 8 days after the cannibalistic bees and naïve bees were in full contact (Experiment C, Fig. 5) or being separated by a wire mesh screen allowing trophallactic contact only (Experiment B, Fig. 3). Development of overt virus infection in a large number of recipient worker bees (Fig. 3c, “DWV-GFP”; Fig. 5c T2 group) was observed. Taken together, these findings suggest that cannibalism combined with trophallaxis allows effective spread of DWV between worker bees (Fig. 6). This is an important finding which showed that even if a small proportion of the workers were actively engaged in cannibalism, the infected tissue could be shared between large numbers of the workers in the colony. Such sharing could increase the impacts of cannibalism on DWV circulation^36^.

**Figure 6.**
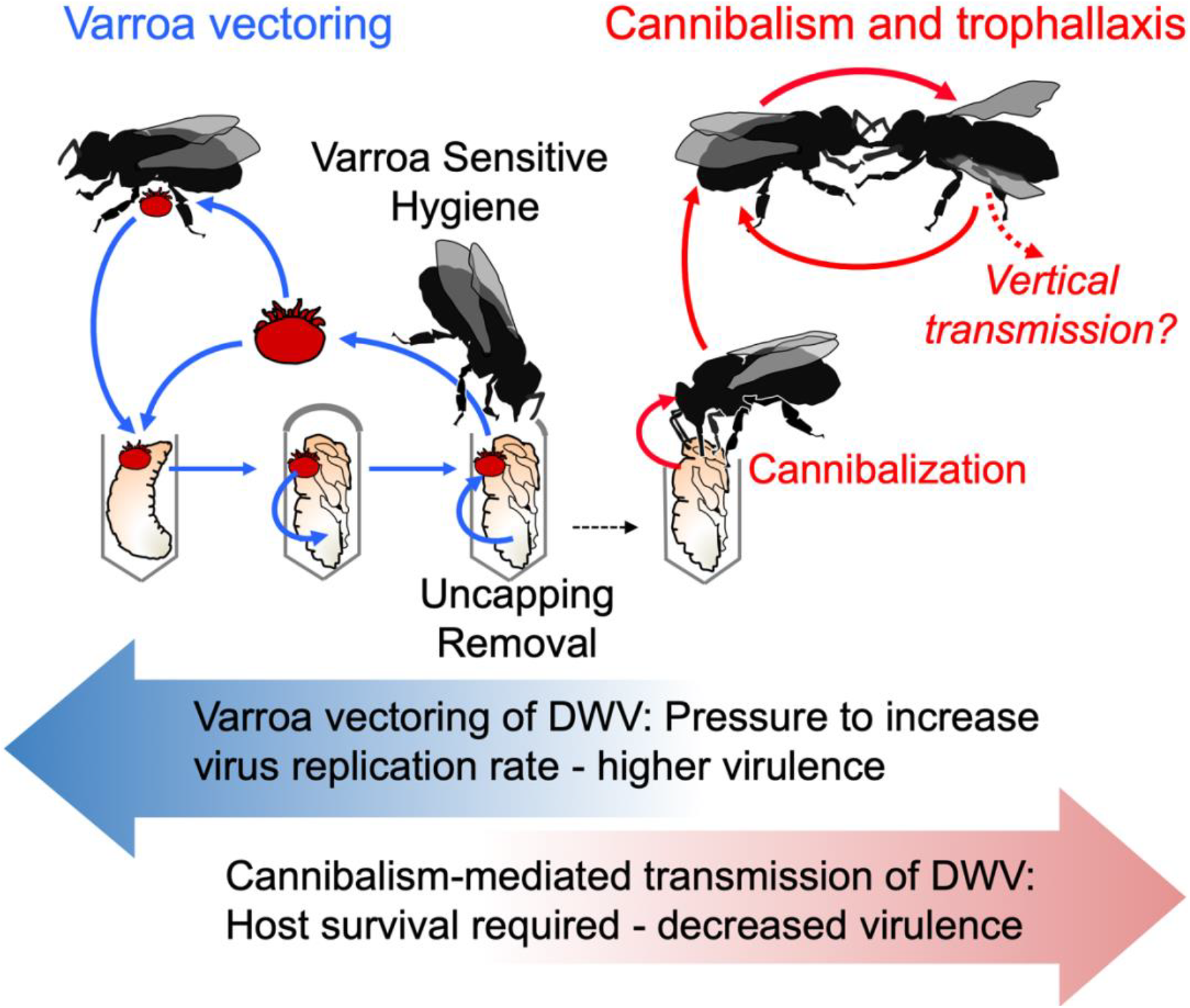
Model of DWV circulation in Varroa mite-infested Varroa Sensitive Hygienic (VSH) colonies. *Varroa* transmission - blue arrows, cannibalism-trophallaxis transmission - red arrows. Block arrows show possible evolutionary pressures which on Varroa and cannibalism-trophallaxis transmission routes impose on DWV virulence.

The use of GFP-tagged DWV gave an additional insight into the mechanisms of trophallactic transmission of DWV. The clone-derived DWV included a non-essential GFP gene, which could be lost from viral genomes during replication^40^, (Fig. 2e). Following replication of this DWV-GFP clone-derived virus population, the proportion of viral genomes with the GFP deletion increased and the loss of GFP could be utilized as a molecular clock. This allowed us to distinguish between the original virus (with nearly 1:1 ration of DWV to GFP copies) and virus populations which had gone through multiple cycles of replication^40^ (Fig. 2e). Therefore, the higher DWV to GFP copy number ratio in worker bees which acquired the virus by trophallaxis from the bees involved in cannibalism (Fig. 4b) suggested transmission of the virus produced after replication events in the worker bees rather than directly from the cannibalized pupal tissues. It is known that hypopharyngeal and mandibular gland secretions of the worker bees could be shared by trophallaxis^26,45^ and DWV was detected in hypopharyngeal glands of worker bees^46^. Efficient transmission and circulation of cannibalism-acquired DWV therefore could depend on survival of the infected bees, thereby selecting against DWV virulence (Fig. 6). Natural attenuation has been reported for RNA viruses, including flaviviruses Japanese encephalitis virus and Dengue virus type-2^47,48^.

Our results suggest that cannibalization of *Varroa*-infested pupae uncapped as a result of VSH activity and trophallactic interactions could provide an efficient route for transmission and circulation of a *Varroa*-vectored DWV (Fig. 6). Oral acquisition of infected pupal tissues with high virus loads and further trophallactic transmission results in DWV infection with the levels typical for overt infection of the virus in the worker bees. This might explain why high DWV loads persist and a poor survival prognosis remains in the colonies which reached a threshold *Varroa* infestation level, even if *Varroa* mites are eliminated *via* varroacide treatments^9^. While VSH is an important trait for reducing mite parasitism, it cannot be excluded that increased VSH activity in *Varroa*-infested hives could lead to increased infection levels and circulation of DWV. Therefore, the cannibalism-trophallactic transmission route of DWV, in addition to *Varroa* vectoring (Fig. 6), should be considered in designing anti-*Varroa* and antivirus treatments of honey bees.

## Methods

### *Honey bees and Varroa* mites

The worker bees used in laboratory cannibalism experiments were sourced in June 2020 from the Beltsville USDA Bee Research Laboratory apiary from a strong colony JC-2 for Experiments A, C, and B (cannibal group), and JC-6 for Experiment B (recipient bees). These colonies had low *Varroa* mite infestation rates (below 0.5 %), and the DWV loads in their pupae were undetectable by qRT-PCR in May and June 2020. To obtain newly emerged workers, the frames from these colonies with sealed brood close to emergence were placed in cages in an environmental chamber set to 32°C and 85% relative humidity in darkness, and newly emerged adult bees were collected after 18 hours incubation. Pupae at the white-eyed stage were pulled out of Varroa-free cells of colony JC-2 using soft tweezers no more than 24 hours prior to their use in the experiments. *Varroa* mites were manually collected from newly emerged drones sourced from additional *Varroa*-infested colonies in the BRL apiary. *Varroa* mites were hand-collected from adult drones from the broodnest of colonies maintained in College Park, MD and the USDA. The colonies had high varroa levels but did not show clinical signs of varroosis. Cannibalism experiments were carried out in dark incubators, at +33°C, relative humidity 85% relative humidity. The worker bees had *ad libitum* access to sugar syrup in a 1:1 ratio accessible and water in the tube feeders changed every 24 hours. For RNA extraction, live bees were sampled and immediately frozen at −80°C. In each experiment there were no significant differences in worker bee mortality between treatment groups.

### Infection of honeybee pupae by DWV-GFP

Honey bee pupae at the white eye developmental stage collected from Varroa-free brood cells were injected with 8 μL of a filtered extract containing 10^7^ of DWV-GFP virus particles. This extract was generated using individual pupae infected with *in vitro* RNA transcript from the construct pDWV-L-GFP carrying an enhanced-GFP coding sequence^40^, which gave an equimolar ratio of DWV to GFP in qRT-PCR tests, indicating that it contained mainly intact recombinant virus without GFP deletions. The extract-injected pupae were incubated in the dark for 48 hr at +33°C, relative humidity 85% prior to development of GFP fluorescence visible when illuminated with long wave, 395 nm, ultraviolet light illumination (Fig. 2a) and were offered for cannibalization (Experiments A and B).

### Analysis of virus replication

Total RNA was extracted from adult honey bee workers or pupae, which were flash-frozen and stored at −80°C. RNA extraction from individual insects included homogenization with 1 mL of Trizol reagent (Ambion) and further purification using RNeasy kits (QIAGEN) according to the manufacturer’s instructions. Extraction of total RNA from pools of frozen worker bees started with lysis in guanidine isothiocyanate buffer as described previously^49^, followed by further disruption using QIAShredder (QIAGEN) and purification using RNeasy kits (QIAGEN). Quantification of DWV and GFP RNA in these RNA extracts was carried out by RT-qPCR as previously^40^ and included cDNA synthesis using Superscript III (Invitrogen) and random hexanucleotides as primers, and qPCR using SYBR green (BioRad) and the primers specific to DWV genomic RNA (5’-GAGATCGAAGCGCATGAACA-3’ and 5’-TGAATTCAGTGTCGCCCATA-3’, positions 6497 - 6626 nt of DWV, positions 7268 – 7397 of DWV-L-GFP), to the region spanning the eGFP - structural VP2 interface (GFP-specific primer 5’-GCATGGACGAGCTGTACAAG-3’, and DWV-specific 5’-CCTTTTCTAATTCAACTTCACC - 3’, positions 2526 - 2624 of DWV-L-GFP genome), and to the honey bee β-actin mRNA (5’-AGGAATGGAAGCTTGCGGTA-3’ and 5’-AATTTTCATGGTGGATGGTGC-3’). The plasmid pDWV-L-GFP^40^ was used as a standard for quantification of DWV and GFP copy numbers, which were log-transformed prior to statistical analyses. One-way analysis of variance (ANOVA) and Tukey’s HSD post-hoc tests were used to assess significance of the differences among the treatment groups.

The cDNA was used to amplify a 1237 nt cDNA fragments corresponding to the 5’ region of DWV RNA (30 to 1266 nt) containing the PacI site introduced into the clone-derived DWV-L-GFP, but absent in the wild-type DWV, using primers 5’-GCCTTCCATAGCGAATTACG-3’ and 5’-CGCCGCCTGGCTTCATCA-3’. The amplicons were digested with *Pac*I restriction enzyme (NEB) for 2 hours, separated by agarose gel electrophoresis and the images were used to estimate the proportion of clone-derived DWV using ImageJ^50^.

## Acknowledgements

This research used resources provided by the SCINet project of the USDA - Agricultural Research Service, ARS project number 0500-00093-001-00-D and was supported by the USDA National Institute of Food and Agriculture grant 2017-06481 to EVR, JDE, and YC. USDA is an equal opportunity provider and employer.

## Author Contributions

F.P.-F., D.J.E and E.V.R., the corresponding authors, conceived the study and contributed equally to this research. Z.L. and D.H. carried out filed monitoring of cannibalism. Y.C. supervised honeybee colony monitoring. F.P.-F, Z.L. and E.V.R. carried work with bees and mites. E.V.R. and F.P.-F carried out the laboratory molecular work and analyzed virus quantification. M.H. analyzed accumulation of the honey bee transcripts. F.P.-F, Z.L., J.D.E. and E.V.R., wrote the manuscript. All co-authors contributed to data interpretation, and to the writing of the manuscript.

---

**Supplementary Table 1.**
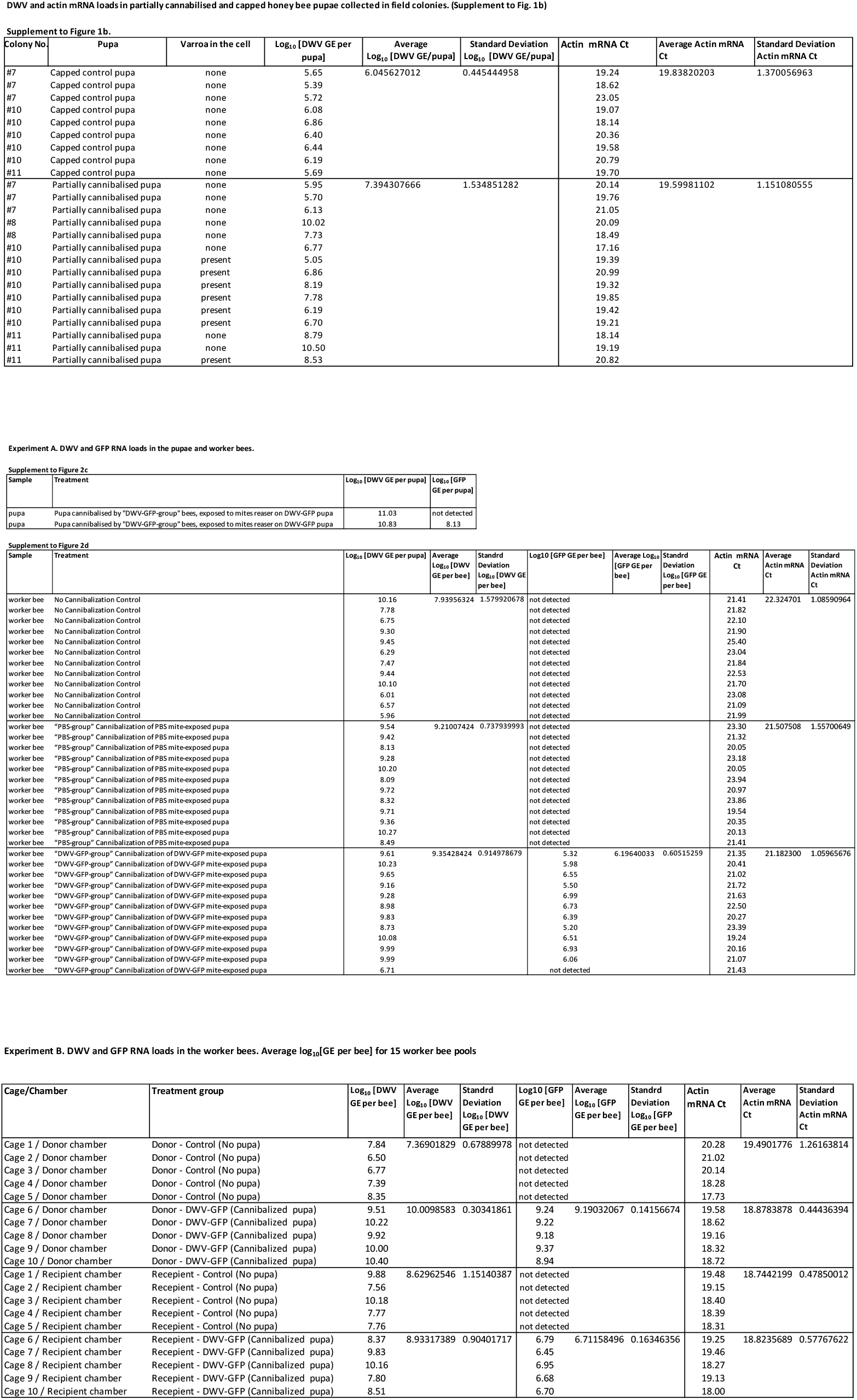
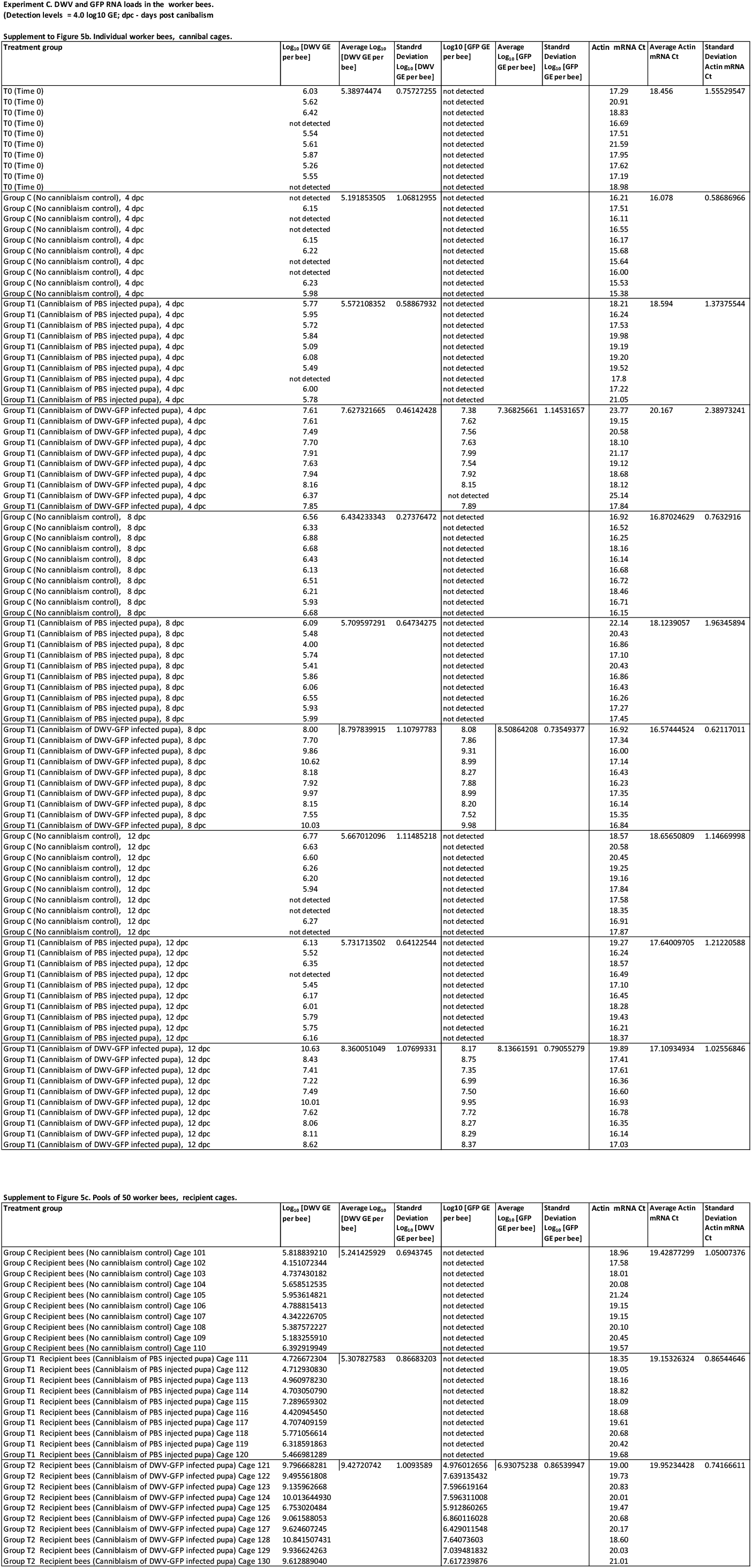

